# Multi-omics Integration with GWAS Unveils Molecular Mechanistic Insights for Type 2 Diabetes

**DOI:** 10.1101/2025.11.21.689764

**Authors:** Malak Abbas, Huishi Toh, Pamela M. Martin, Merry L. Lindsey, Mileati T. Melese, Antwi-Boasiako Oteng, Amadou Gaye

**Author notes:** Corresponding author: Amadou Gaye, PhD, 1005 Dr DB Todd Jr Blvd, Nashville, TN 37208.

## Abstract

**Background:** Type 2 diabetes (T2D) is a complex metabolic disorder driven by genetic and environmental factors. While genome-wide association studies (GWAS) have identified numerous T2D-associated variants, many remain functionally uncharacterized. Integration of GWAS with molecular phenotyping offers a path to revealing biological relevance. We investigated the influence of GWAS-variants, including sub-threshold T2D-associated variants (GWAS p-value ≤ 0.0001), on gene and protein expression to assign functional relevance.

**Methods:** Genetic variants associated with T2D in the GWAS Catalog and present in our whole-genome sequencing (WGS) data were used to perform expression quantitative trait loci (eQTL) analysis in 242 whole-blood mRNA-sequenced samples. The same variants were used to perform protein quantitative trait loci (pQTL) analysis in a set of 362 plasma samples profiled on the Olink platform. For each analysis, the datasets were randomly split into discovery and validation subsets. Associations between variants and mRNA or protein levels were tested by multiple linear regression, and only QTLs that reached a false discovery rate adjusted p-value ≤ 0.05 in the discovery dataset and replicated in the validation dataset (p ≤ 0.05) with same direction of effect were carried forward. QTL-linked mRNAs and proteins were subsequently evaluated for their relationship with T2D status to connect them with T2D pathophysiology.

**Results:** We identified 1,291 eQTLs linked to 97 mRNAs and 1,273 pQTLs linked to 22 proteins. Among these, 10 mRNAs and 5 proteins were differentially expressed between non-diabetic and diabetic individuals. Notably, LPL, APOBR, APOM (lipid metabolism), NOTCH2, TREH (β-cell/endocrine regulation), and HLA-A, OAS3 (immune response) converged on three biological axes central to T2D pathophysiology. The directionality of molecular effects was consistent with known disease mechanisms, including insulin resistance (LPL, APOBR), β-cell stress (TREH, NOTCH2), and chronic inflammation (OAS3).

**Conclusions:** Our findings indicate that variants falling below conventional GWAS significance thresholds can have demonstrable effects on gene expression and protein levels. This underscores the importance of prioritizing biological relevance alongside statistical significance, rather than relying solely on rigid p-value cutoffs.

## INTRODUCTION

Type 2 diabetes (T2D) imposes a significant public health burden, particularly in the United States, where its prevalence has surged to epidemic proportions. Recent data from the Centers for Disease Control and Prevention (CDC) indicates that approximately 34.2 million individuals in the US, constituting roughly 10.5% of the population, are affected by diabetes, with the overwhelming majority (approximately 90-95%) having T2D ^1^. We focus on individuals of African American (AA) ancestry, a population that exhibits a disproportionately higher prevalence of T2D and an elevated risk of diabetes-related complications ^2^. Investigating this group is critical for elucidating the biological, environmental, and social determinants contributing to these disparities and for informing targeted interventions aimed at reducing the burden of T2D across all communities. Genetic predisposition contributes significantly to T2D susceptibility; for instance, carriers of the TCF7L2 risk allele (rs7903146) have a ∼1.5-fold higher risk of T2D.

Genome-wide association studies (GWAS) have identified dozens of genetic variants associated with T2D risk ^3–10^. Translating GWAS findings of T2D into clinical advances, however, presents notable challenges. One key obstacle is the sheer complexity of the disease, as T2D is influenced by numerous genetic variants with most exerting a modest effect size ^11^. This intricate genetic landscape makes it challenging to pinpoint more precisely genes and pathways driving T2D pathogenesis.

Examining the impact of GWAS variants on gene expression (transcriptome) and protein abundance (proteome) can remove this challenge by coupling functional significance to variant expression. For example, identifying GWAS variants that alter gene expression or protein function can provide mechanistic insights into disease biology ^12^. Furthermore, by integrating GWAS data with transcriptomic and proteomic datasets, we can link causal genes and pathways to underlying disease associations ^13^. For instance, GWAS variants may regulate gene expression levels, leading to downstream effects on biological pathways implicated in disease pathogenesis ^14^. Integrating GWAS data with transcriptomic and proteomic data enables a comprehensive understanding of disease biology at the systems level.

We investigated the effect of genetic variants associated with T2D in GWAS on gene and protein expression in a cohort of AA subjects. We tested the hypothesis that GWAS variants associated with concomitant changes in gene or protein expression would indicate functional variants including some below the conventional GWAS statistical significance threshold.

## MATERIAL AND METHODS

### Data Source

The GENomics, Environmental FactORs and the Social DEterminants of Cardiovascular Disease in African Americans STudy (GENE-FORECAST) is a research platform strategically designed to employ a comprehensive, multi-omics systems biology approach for in-depth, multi-dimensional phenotyping of health and disease within the AA population. Utilizing a community-based sampling framework, GENE-FORECAST established a cohort comprising U.S.-born AA men and women aged 21-65, recruited primarily from the metropolitan Washington D.C. area as described previously ^15^. GENE-FORECAST as approved by the Institutional Review Board (IRB) of the National Institutes of Health and was conducted in compliance with local regulations and institutional protocols. All participants provided written informed consent prior to their involvement. The data utilized in this project are from GENE-FORECAST.

### Clinical Features

Clinical information was collected to assess type 2 diabetes (T2D) status and related metabolic traits. Fasting blood glucose (FBG), HbA1c, and homeostatic model assessment of insulin resistance (HOMA-IR) were measured in all participants. A total of 664 subjects were assessed and classified as diabetic (n = 79) if they self-reported a physician diagnosis or reported current use of T2D medication, or as non-diabetic (n = 585). The baseline characteristics of the 664 individuals are reported in Table 1.

**Table 1:**
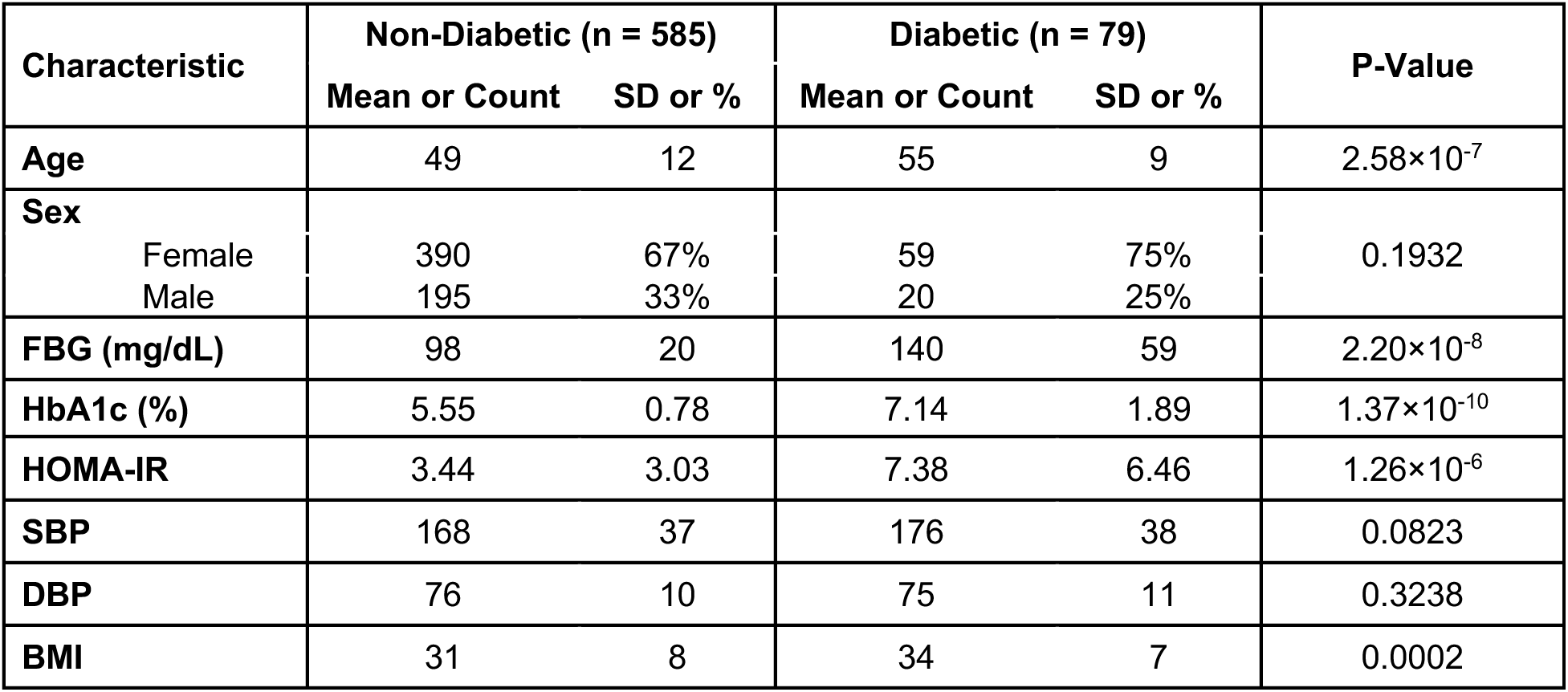
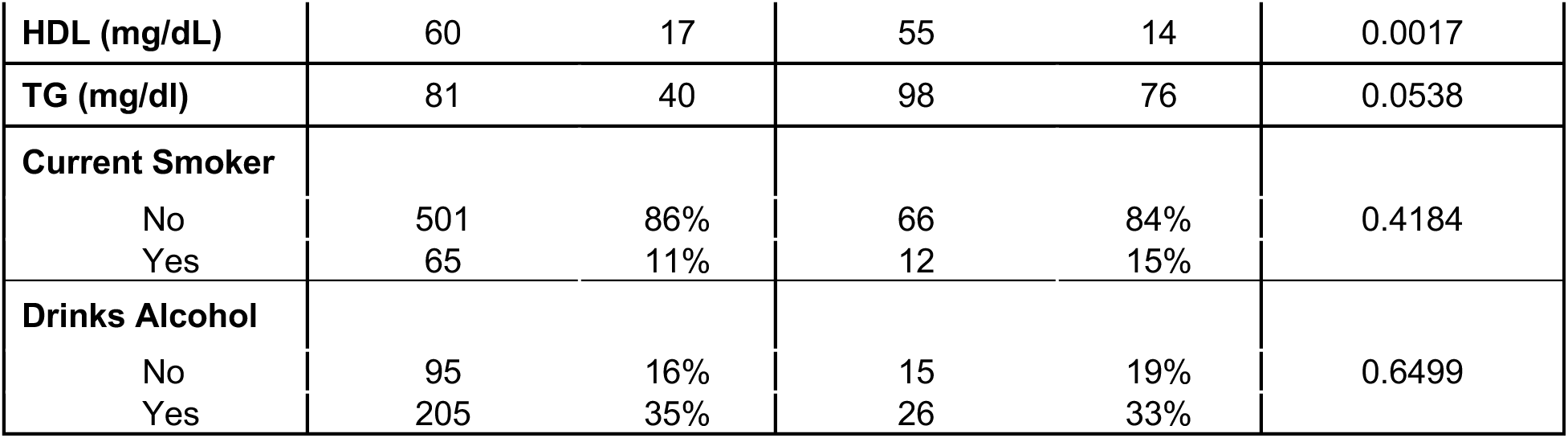
Baseline clinical and demographic characteristics of study participants, stratified by type 2 diabetes (T2D) status. Values are shown as mean ± standard deviation (SD) for continuous variables and count and percentage for categorical variables. P-values correspond to group comparisons between non-diabetic (n = 585) and diabetic (n = 79) individuals.

For subsequent multi-omics analyses, only subsets of these 664 individuals had data available for specific modalities (see Figure 1). Whole-genome sequencing (WGS) was available for 371 subjects, whole-blood mRNA sequencing (transcriptome) for 497, and plasma protein levels (proteome) for 638. Because some assays were conducted at later stages as new participants were enrolled and given the high cost and resource constraints associated with certain platforms, not all individuals had data available across every modality, resulting in differing sample sizes for each analysis. Accordingly, expression quantitative trait loci (eQTL) analyses were conducted on 242 individuals with both transcriptome and WGS data, while protein quantitative trait loci (pQTL) analyses were performed on 362 individuals with both proteome and WGS data.

**Figure 1:**
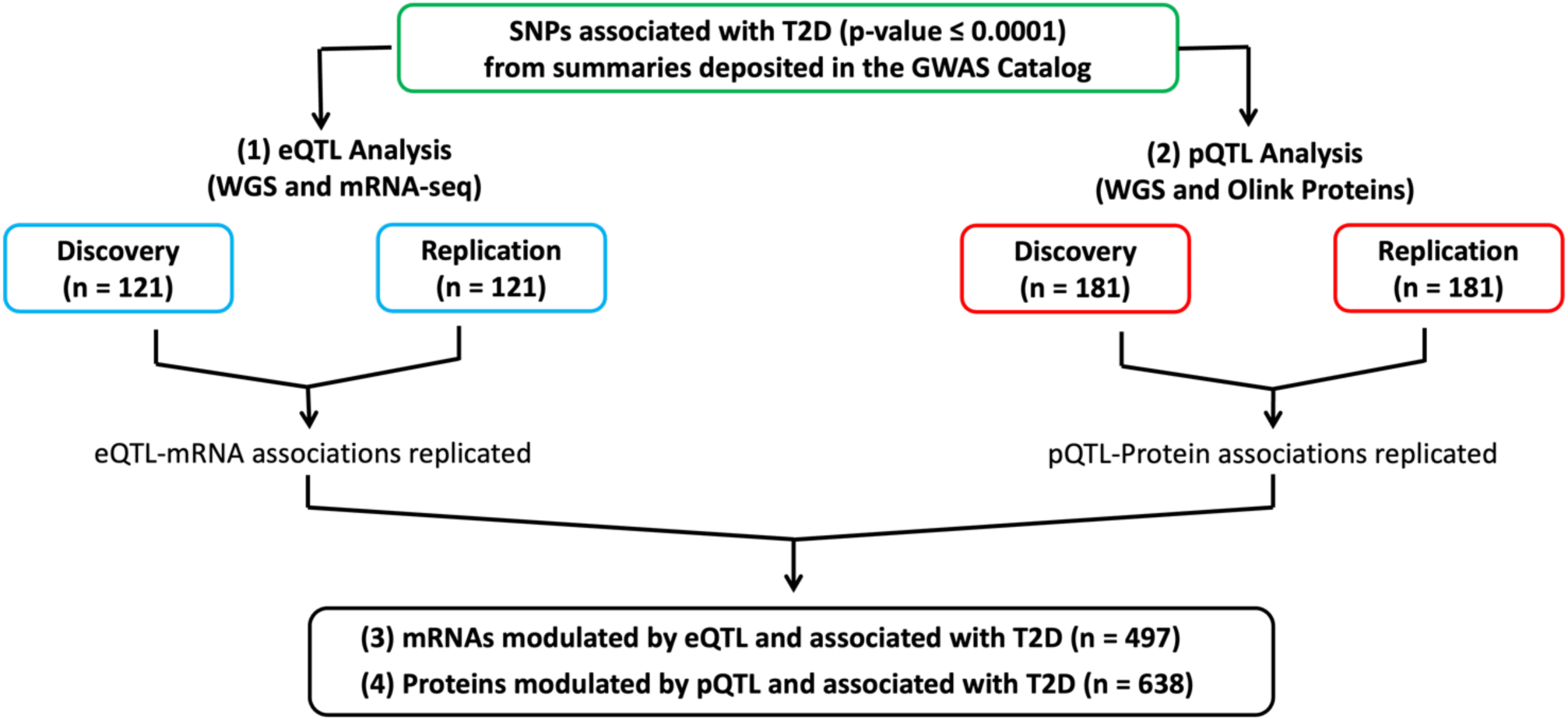
Overview of the analyses performed to identify (1) eQTLs associated with mRNAs in both datasets, (2) pQTLs associated with proteins in both datasets, (3) eQTL-linked mRNAs associated with T2D and (4) pQTL-linked proteins associated with T2D.

### Genotype Data

The genotype data are from WGS. DNA was extracted from whole blood collected in ethylenediaminetetraacetic acid (EDTA) coated tubes followed by picoGreen quantitation. Library preparation was done using Whole Genome Small Insert PCR-Free. The WGS samples were 151bp paired end sequenced on an Illumina NovaSeq6000 to a mean coverage of 30X. At the data preprocessing step, WGS reads were processed with the Whole Genome Germline Variant Discovery pipeline developed and used by the Genomics Platform at the Broad Institute. Reads were aligned to the genome build Hg38 with Burrows-Wheeler Aligner and gVCF generation, joint genotype calling, and quality filtering were executed in accordance with GATK4 best-practices. Of the 371 samples with whole-genome sequencing (WGS) data, 242 also had whole-blood gene expression profiles obtained through mRNA sequencing, and 362 had plasma protein levels measured using the Olink Explore 3072 platform.

### Transcriptomics Data

The transcriptome data consists of the messenger RNA sequencing (mRNA-seq) from 497 whole blood samples. Total RNA extraction was carried out using MagMAXTM for Stabilized Blood Tubes RNA Isolation Kit (Life Technologies, Carlsbad, CA). For the library preparation, total RNA samples were converted into indexed cDNA sequencing libraries using Illumina’s TrueSeq kits. Ribosomal RNA (rRNA) was removed. The GENE-FORECAST samples were pair-end sequenced on the Illumina HiSeq2500 and HiSeq4000 platforms with a sequencing depth of at least 50 million read per sample. To measure gene expression levels, mRNA expression was quantified using a bioinformatics pipeline developed by the Broad Institutes and used by the Genotype-Tissue Expression (GTEx). The pipeline is detailed in the GitHub software development platform ^16^. Transcripts with expression levels below 2 counts per million (CPM) in at least 90% of samples were excluded from the analysis. Out of 55,010 total mRNA transcripts, 18,871 did not meet this criterion and were removed. The expression data was normalized using the Trimmed Mean of M-values (TMM), an optimal method for read count data ^17^. Principal component analysis (PCA) was conducted to identify and exclude transcript outliers; 4 transcripts were removed. A total of 17,947 protein coding mRNA were included in the expression quantitative trait loci (eQTL) across a subset of 242 samples.

### Proteomics Data

The Explore 3072 assay, utilizing 8 Explore 384 Olink panels (Cardiometabolic, Cardiometabolic II, Inflammation, Inflammation II, Neurology, Neurology II, Oncology, Oncology II), was used to assess the relative expression of a total of 2,947 proteins. Proximity Extension Assay (PEA) technology was conducted according to the Olink AB manufacturer procedures by the certified laboratory. Briefly, the technique relies on the use of antibodies labelled with unique DNA oligonucleotides that bind to their target protein present in the sample. The DNA oligonucleotides, when in proximity of a target protein, undergo hybridization and act as a template for DNA polymerase-dependent extension, forming a unique double-stranded DNA barcode proportionate to the initial protein concentration. Quantification of resulting DNA amplicons is accomplished through high-throughput DNA sequencing, generating a digital signal reflective of the number of DNA hybridization events corresponding to the protein concentration in the original sample. The measurement of protein levels is based on Normalized Protein eXpression (NPX) values, serving as a relative protein quantification unit. This quantification is normalized to account for systematic noise arising from sample processing and technical variation, leveraging internal controls and sample controls. NPX units are on a log2 scale, where one NPX unit increase indicates a two-fold rise in the concentration of amplicons representing the target protein compared to the internal control.

EDTA-derived plasma samples of 635 subjects from the GENE-FORECAST were sent to the Olink Proteomics Analysis Service in Boston, USA. Proteomic analyses were conducted collectively in a single batch. A total of 2,941 proteins that passed QC filters were considered for the protein quantitative trait loci (pQTL) analysis across a subset 362 samples.

### Statistical Analyses

The GENE-FORECAST data were randomly split into two datasets: the discovery dataset and the validation dataset. The analyses were conducted in three main steps depicted graphically in Figure 1. First, summaries from GWAS studies of T2D deposited in the GWAS catalog were downloaded. Single-nucleotide polymorphisms (SNPs) reported as associated with T2D at a p-value ≤ 0.0001 were extracted. Then, eQTL and pQTL analyses were conducted with that set of SNPs. Finally, the relationship between genes and proteins linked to the eQTL and pQTL variants was investigated using the whole GENE-FORECAST dataset.

#### – SNPs associated with mRNA expression (eQTL) and protein level (pQTL)

eQTL and pQTL analyses were performed to assess the impact of SNPs previously associated with T2D in GWAS on gene expression and protein abundance, thereby providing insights into the molecular mechanisms linking GWAS-identified variants to T2D pathophysiology.

Both eQTL and pQTL analyses were conducted utilizing the *MatrixEQTL* ^18^ R library. *MatrixEQTL* applies a regression model with mRNA/protein levels as the outcome and additive genotypes as independent variables. The regression model was adjusted for covariates, including age, sex, and principal components (PCs) 1 to 6 that explain most of the variance, to account for genetic ancestry admixture. The analyses were restricted to variants located within the cis region, spanning 1 megabase, of the mRNA for eQTL or the mRNA encoding the protein for pQTL.

A SNP was designated as an eQTL if it met all three criteria: (a) the multiple testing-adjusted p-value in the discovery set was equal to or less than 0.05; (b) the nominal p-value in the validation dataset is equal to or less than 0.05; and (c) the beta value of the association with an mRNA was in the same direction in the discovery and validation datasets. Analogously, a SNP was deemed a pQTL if the beta value and the p-value of the association with a protein meet the same three criteria.

The focus on bi-allelic SNPs for the analyses was grounded on multiple considerations, including the higher accuracy and reliability associated with genotyping techniques for bi-allelic SNPs compared to multi-allelic SNPs. This increased precision contributes to the robustness of our pQTL and eQTL findings. Furthermore, the choice of bi-allelic SNPs is strategically driven by their binary allele composition, facilitating the interpretation of genetic effects and streamlining the identification and characterization of associations between specific alleles and mRNA and protein levels.

#### – eQTL/pQTL-linked mRNAs/proteins associated with T2D

The relationship between T2D and mRNAs and proteins linked to QTLs was evaluated. A differential expression (DE) analysis was carried out with the list of mRNAs associated with eQTLs, in a sample of 497 individuals (53 diabetic and 444 non-diabetic) for whom mRNA-seq data and T2D status was available. To conduct the DE analysis, we utilized the R library *edgeR* which fits a negative binomial model to mRNAs read counts and computes likelihood ratio tests for the coefficients in the model. The model was adjusted for age and sex. An mRNA was considered significantly differentially expressed between diabetic and non-diabetic subjects if the Benjamini-Hochberg (BH) false discovery rate (FDR) adjusted p-value ≤ 0.05.

The association between proteins linked to pQTLs and T2D was investigated by fitting a generalized linear model (GLM) with the protein level as outcome variable and T2D as exposure variable. The sample size for the protein-T2D association analysis consists of 635 subjects (73 diabetic and 562 non-diabetic) for whom protein data and T2D was available. The model was adjusted for age and sex and a protein was considered significantly associated with T2D if the BH’s FDR adjusted p-value was equal to or less than 0.05.

## RESULTS

### Baseline Data Analysis

Table 1 presents baseline characteristics of 664 participants (585 non-diabetic, 79 diabetic). Diabetic individuals were older (55 ± 9 years) compared with non-diabetic participants (49 ± 12 years; P = 2.58×10^-7^) and showed higher fasting blood glucose (176 ± 38 vs. 168 ± 37; P = 2.20×10^-8^), HbA1c (7.14 ± 1.89 vs. 5.55 ± 0.78; P = 1.37×10^-10^), and HOMA-IR (7.38 ± 6.46 vs. 3.44 ± 3.03; P = 1.26×10^-6^). BMI was also significantly elevated among diabetics (34 ± 7 vs. 31 ± 8; P = 0.0002) whilst HDL levels were lower in diabetics (55 ± 14 vs. 60 ± 17; P = 0.0017). No significant differences were observed in sex distribution, blood pressure (SBP and DBP), current smoking status, or alcohol consumption.

### GWAS studies of T2D extracted from the GWAS Catalog

The summary statistics of seven GWAS studies of T2D studies were downloaded from the GWAS Catalog ^19^ (version of December 2023) and all variants with a significance level p-value ≤ 0.0001 were extracted. The seven GWAS studies of T2D are presented in Table 2.

**Table 2:** GWAS studies of T2D with summary statistics available from the GWAS catalogue.

| GWAS Catalog Study Accession | PubMed ID | Publication Date (Author) | Study Title | Study Sample Description |
| --- | --- | --- | --- | --- |
| GCST90255648 | 37148359 | 2023-05-06 (Huerta-Chagoya et al.) | The power of TOPMed imputation for the discovery of Latino-enriched rare variants associated with type 2 diabetes | 8,150 Hispanic or Latin American cases, 10,735 Hispanic or Latin American controls |
| GCST90161239 | 36329257 | 2022-11-03 (Lee et al.) | Phenome-wide analysis of Taiwan Biobank reveals novel glycemia-related loci and genetic risks for diabetes | 3,844 Taiwanese ancestry cases, 59,333 Taiwanese ancestry controls |
| GCST90093109 | 35393509 | 2022-04-07<br>(Loh et al.) | Identification of genetic effects underlying type 2 diabetes in South Asian and European populations | 16,677 South Asian ancestry cases, 33,856 South Asian ancestry controls |
| GCST90043638 | 34737426 | 2021-11-04<br>(Jiang et al.) | A generalized linear mixed model association tool for biobank-scale data | 455,634 European ancestry controls |
| GCST90081516 | 34662886 | 2021-10-18<br>(Backman et al.) | Exome sequencing and analysis of 454,787 UK Biobank participants. | 30,259 European ancestry cases, 355,886 European ancestry controls |
| GCST90018706 | 34594039 | 2021-09-30<br>(Sakaue et al.) | A cross-population atlas of genetic associations for 220 human phenotypes | 45,383 East Asian ancestry cases, 132,032 East Asian ancestry controls |
| GCST007515 | 29632382 | 2018-04-09<br>(Mahajan et al.) | Refining the accuracy of validated target identification through coding variant fine mapping in type 2 diabetes | 48,286 European ancestry cases, 250,671 European ancestry controls, 33,126 African American, East Asian, Hispanic/Latino or South Asian cases, 120,161 African American, East Asian, Hispanic/Latino or South Asian controls |

Subsequently, genomic positions on the Genome Reference Consortium Human Build 38 (GRCh38) were used to evaluate the intersection between the GWAS variants and bi-allelic SNPs in the GENE-FORECAST WAGS that have a minor allele frequency (MAF) of 0.005 or more. Among the variants extracted from the GWAS summaries, a total of 34,176 bi-allelic SNPs were present in the GENE-FORECAST WGS.

### SNP-mRNA associations

The SNP-mRNA association analysis included 34,176 SNPs and 89,147 mRNA transcript isoforms from 17,947 protein-coding genes, within both the GENE-FORECAST discovery and validated datasets. Following eQTL analysis, 3,014 statistically significant SNP-mRNA associations (adjusted p-value ≤ 0.05) were identified within the discovery dataset with a nominal p-value ≤ 0.05 and subsequently confirmed in the replication dataset. These replicated eQTL-mRNA associations comprised 1,291 distinct eQTLs and 97 unique mRNA transcripts. Most of the eQTLs, 90% are common (MAF ≥ 0.05), in the GENE-FORECAST dataset. Details of the replicated SNP-mRNA associations are provided in Supplemental Table T1. A graphical summary of the results is provided in Figure 2.

**Figure 2:**
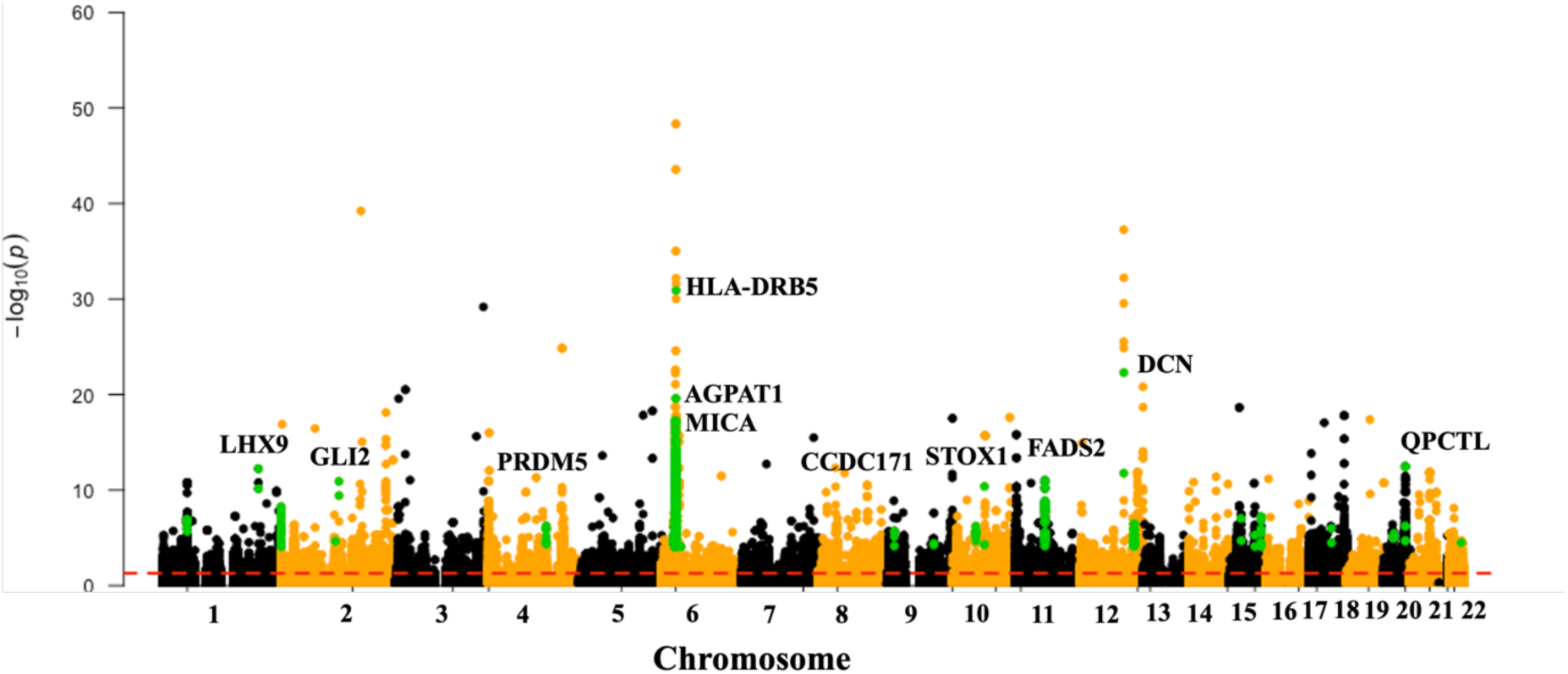
Summary of the SNP-mRNA associations in GENE-FORECAST. SNPs in validated associations are depicted in green color. The black and orange band represent the chromosomes and SNPs tested within each chromosome.The dashed red line is the threshold for statistical significance (FDR adjusted p-value ≤ 0.05). The mRNAs in the validated associations with the lowest p-value are labelled.

### eQTL-linked mRNA differentially expressed by T2D status

Among the 97 mRNAs associated with eQTLs, 10 (10%) were differentially expression by T2D in a sample set comprising 497 subjects (53 diabetic and 444 non-diabetic) from the GENE-FORECAST study. Table 3 provides an overview of these 10 mRNAs, detailing their differential expression patterns and the corresponding count of unique eQTLs linked to each mRNA within replicated eQTL-mRNA associations. Notably, 8 of the 10 mRNAs (80%) were downregulated in the diabetic group. Further insights into the potential genetic underpinnings of these mRNA-T2D associations are presented in Table 4, which lists eQTL-mRNA pairs, with the top SNP associated with each of the 10 mRNAs based on its GWAS p-value. Additionally, the effect size and significance level of the association between the SNP and T2D in the GWAS study are provided.

**Table 3:** mRNA differentially expressed by T2D status and associated with one or more eQTL. The table provides the log fold-change (logFC), p-value and FDR adjusted p-value of the differential expression analysis. The last column provides the number of eQTL associated with the mRNA in the replicated eQTL findings.

| Transcript | mRNA Symbol | Gene Name | Chr | logFC | P | FDR P | # eQTL associated |
| --- | --- | --- | --- | --- | --- | --- | --- |
| ENST00000473954 | *PPP1R10* | Protein Phosphatase 1 Regulatory Subunit 10 | 6 | -2.17 | 1.48e-07 | 2.26e-05 | 1 |
| ENST00000523235 | *FADS2* | Fatty Acid Desaturase 2 | 11 | -1.16 | 4.21e-05 | 0.002 | 27 |
| ENST00000361492 | *GLI2* | GLI Family Zinc Finger 2 | 2 | -1.75 | 4.50e-05 | 0.002 | 1 |
| ENST00000680435 | *CTSV* | Cathepsin V | 9 | -1.81 | 2.06e-04 | 0.008 | 1 |
| ENST00000259455 | *GABBR2* | Gamma-Aminobutyric Acid Type B Receptor | 9 | -1.59 | 2.79e-04 | 0.010 | 1 |
| ENST00000355484 | *FADS2* | Fatty Acid Desaturase 2 | 11 | -1.43 | 3.75e-04 | 0.013 | 3 |
| ENST00000615520 | *LHX9* | LIM Homeobox 9 | 1 | -1.61 | 8.12e-04 | 0.022 | 1 |
| ENST00000522359 | *FADS2* | Fatty Acid Desaturase 2 | 11 | -1.23 | 9.53e-04 | 0.025 | 19 |
| ENST00000629380 | *SYNGAP1* | Synaptic Ras GTPase Activating Protein 1 | 6 | -1.32 | 1.09e-03 | 0.027 | 1 |
| ENST00000681594 | *OAS3* | 2'-5'-Oligoadenylate Synthetase 3 | 12 | -0.76 | 2.13e-03 | 0.044 | 5 |

**Table 4:**
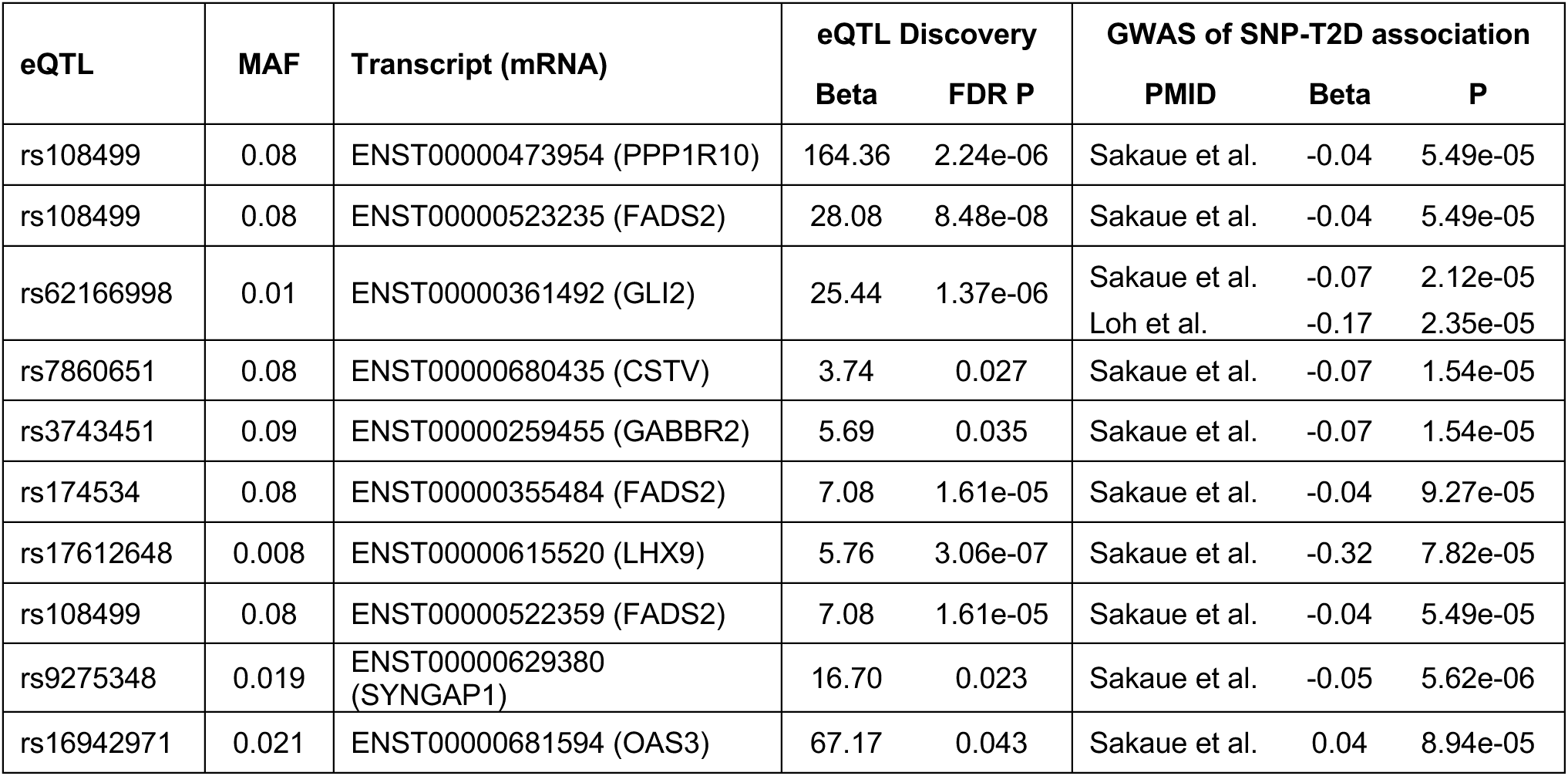
Summary of SNP-mRNA association results for the 10 transcripts differentially expressed by T2D. If a transcript is associated with multiple SNPs, only the top SNP by GWAS p-value is included in the table.

### SNP-Protein associations

A total of 1,324 statistically significant associations SNP-protein associations were observed in the discovery dataset and replicated; those associations involved 1,273 unique pQTLs and 22 unique proteins. Most of the pQTL (97%) are common (MAF ≥ 0.05). Details of the replicated pQTL-protein associations are available in Supplemental Table T1. A graphical summary is provided in Figure 3.

**Figure 3:**
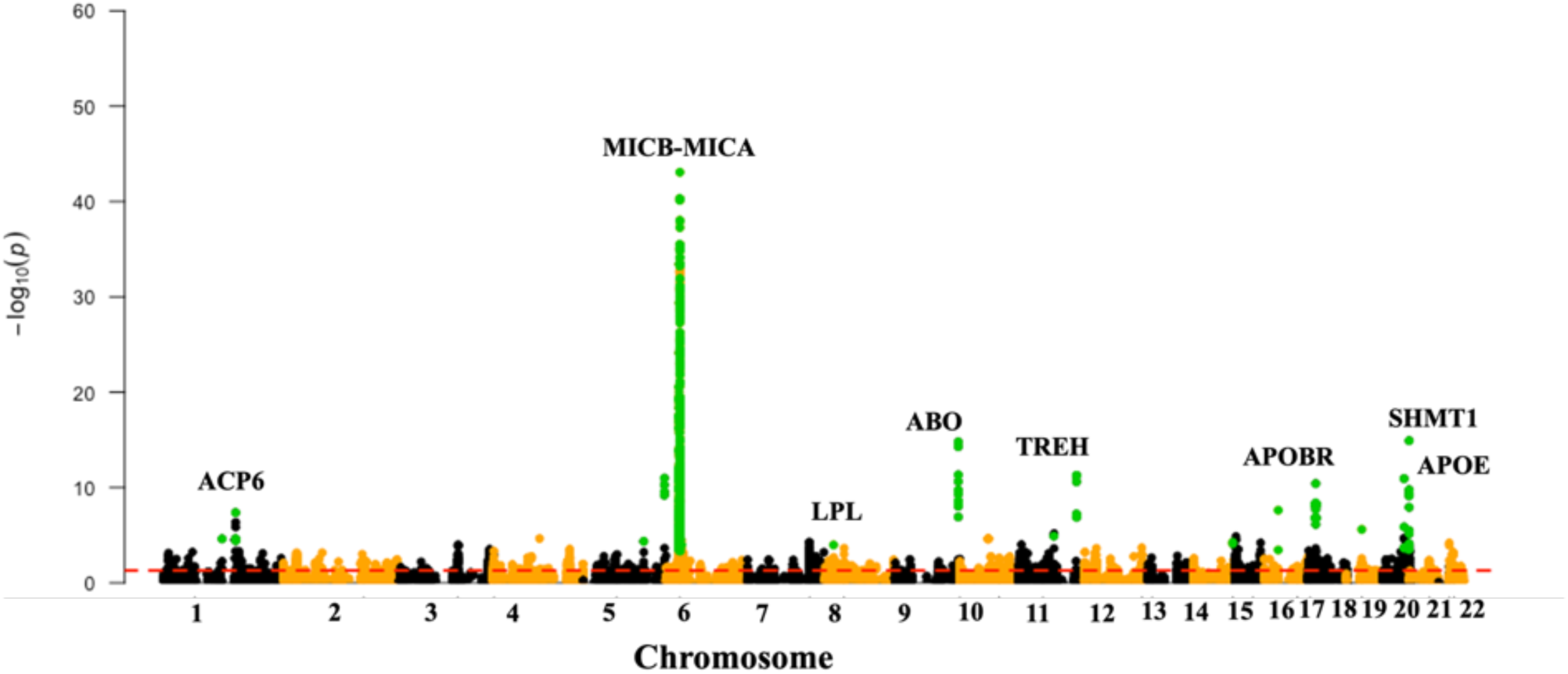
Summary of SNP-protein associations identified in GENE-FORECAST. Validated associations are highlighted in green, while black and orange bands represent chromosomes and SNPs tested per chromosome. The red dashed line denotes the statistical significance threshold (FDR-adjusted p-value ≤ 0.05). Proteins with the most significant associations are labeled.

### pQTL-linked proteins associated with T2D

Among the 22 proteins linked to pQTLs, five (23%) exhibited associations with T2D within a sample set comprising 635 subjects (73 diabetic and 562 non-diabetic) from the GENE-FORECAST study. Table 5 delineates these five proteins associated with T2D, along with the respective count of unique pQTLs linked to each protein within replicated pQTL-mRNA associations. T2D was associated with decreased levels of four proteins (APOM, APOBR, LPL, and NOTCH2), while exhibiting an opposing effect for one protein (TREH). Table 6 supplements these findings by presenting pQTL-Protein pairs with the top SNP associated with each of the five proteins based on its GWAS p-value, alongside the effect size and p-value of the association between the SNP and T2D in the GWAS study.

**Table 5:** Proteins associated with T2D status and linked to one or more pQTL. The table provides the beta value, p-value and FDR adjusted p-value of the protein-T2D association. The last column provides the number of pQTL associated with the pQTL in the replicated pQTL findings.

| Protein | Name | Chr. | Beta | P | FDR P | # pQTL associated |
| --- | --- | --- | --- | --- | --- | --- |
| APOM | Apolipoprotein M | 6 | -0.17 | 3.50E-04 | 0.008 | 1 |
| APOBR | Apolipoprotein B Receptor | 16 | -0.19 | 2.12E-03 | 0.023 | 2 |
| LPL | Lipoprotein Lipase | 8 | -0.26 | 5.57E-03 | 0.041 | 1 |
| TREH | Trehalase | 11 | 0.34 | 9.23E-03 | 0.051 | 6 |
| NOTCH2 | Notch Receptor 2 | 1 | -0.12 | 1.50E-02 | 0.066 | 1 |

**Table 6:** Summary of SNP-Protein association results for the 5 proteins associated with T2D. For the proteins associated with multiple SNPs, only the top SNP by GWAS p-value is included in the table.

| pQTL | MAF | Protein | pQTL Discovery |  | GWAS of SNP-T2D association |  |  |
| --- | --- | --- | --- | --- | --- | --- | --- |
|  |  |  | Beta | FDR P | PMID | Beta | P |
| rs760293 | 0.25 | APOM | -0.20 | 8.29E-03 | Mahajan et al. | -0.04 | 3.37E-05 |
| rs8049439 | 0.43 | APOBR | -0.17 | 4.06E-02 | Mahajan et al. | -0.03 | 2.02E-05 |
| rs7016880 | 0.05 | LPL | -0.64 | 1.47E-02 | Mahajan et al. | -0.05 | 1.05E-06 |
| rs10892252 | 0.07 | TREH | -1.11 | 2.35E-05 | Sakaue et al. | -0.05 | 1.36E-05 |
| rs835574 | 0.18 | NOTCH2 | -0.25 | 3.80E-03 | Mahajan et al. | 0.04 | 2.91E-06 |

## DISCUSSION

A wide range of genetic factors contribute to the architecture of T2D, influencing lipid metabolism, immune regulation, pancreatic endocrine function, and glucose homeostasis^20,21^. Through multi-omic integration of GWAS loci with transcriptomic and proteomic profiles, we defined functional links between T2D-associated SNPs and molecular traits that provide mechanistic insight into the disease. A major strength of this study is the integration of multi-omics datasets, which allows for a more comprehensive exploration of genetic contributions to T2D. The use of a well-characterized African American cohort also addresses a critical gap in genetic research, the limited diversity of human populations.

Together, our findings define three interrelated axes of lipid metabolism (LPL, APOBR, APOM), endocrine regulation (NOTCH2, TREH), and immune activation (HLA-A, OAS3), through which genetic risk variants exert coordinated effects on molecular traits and disease phenotype. The concordance between genetic associations and differential expression in T2D cases strengthens causal inference and supports these loci as mechanistically relevant contributors to disease.

Among the most prominent signals were loci impacting lipid metabolism. For instance, the SNP rs7016880 within the LPL locus was associated with reduced lipoprotein lipase (LPL) protein level, a pattern similarly observed in individuals with T2D. LPL is a critical enzyme in triglyceride hydrolysis; its downregulation may impair fatty acid uptake and promote ectopic lipid deposition in insulin-responsive tissues, exacerbating lipotoxicity and insulin resistance^22,23^. This aligns with previous studies linking LPL deficiency and certain genetic polymorphisms to elevated plasma triglycerides and insulin resistance, while the S447X variant of LPL, which enhances enzymatic activity, is associated with improved insulin sensitivity and a reduced risk of T2D^24–26^.

Similarly, rs8049439 in APOBR was linked to reduced apolipoprotein B receptor levels, which may impair lipid clearance and promote inflammation^27^. APOBR plays a role in hepatic and macrophage uptake of triglyceride-rich remnants^28,29^; its deficiency may lead to impaired lipid clearance and promote systemic inflammation^30,31^. At the APOM locus, rs760293 was associated with diminished levels of apolipoprotein M, a key S1P-binding protein involved in vascular protection and anti-inflammatory signaling^32,33^, further implicating disrupted lipid signaling in disease progression. The concomitant decline in APOM observed in T2D patients suggests a loss of protective S1P signaling^34^, potentially contributing to vascular and metabolic dysfunction^35^. These functional associations are supported by prior research, which links the dysregulation of LPL, APOBR, and APOM to impaired lipid metabolism, inflammation, and insulin resistance in T2D^36–38^. Collectively, the variants identified converge on the impairment of lipid clearance and signaling, reinforcing their relevance to insulin resistance and metabolic inflammation.

Beyond metabolic regulation, our findings also highlighted genetic influences. In our study, rs835574 in NOTCH2 was associated with reduced Notch2 protein levels in genotype-stratified pQTL analysis, and Notch2 levels were also lower in individuals with T2D. Given the essential role of Notch2 in pancreatic β-cell development and maintenance^39,40^, this suggests that the risk allele may compromise β-cell identity and regenerative capacity. Additionally, rs10892252 near TREH was associated with increased trehalase protein expression, a pattern recapitulated in T2D samples. TREH hydrolyzes dietary trehalose into glucose, and its upregulation may enhance intestinal glucose absorption^41,42^, thereby contributing to postprandial hyperglycemia and exacerbating β-cell stress^43^. Consistent with our findings, prior studies have linked altered NOTCH2 signaling with impaired β-cell development and glucose intolerance^44,45^, while increased TREH activity has been associated with elevated glucose absorption and disrupted glycemic control^46,47^.

Immune-related mechanisms also emerged. rs2517672 was associated with elevated expression of HLA-A, a class I MHC molecule involved in antigen presentation^48^. This observation is consistent with what is known about T2D and suggests a role in aberrant T-cell activation and chronic inflammation. Furthermore, rs16942971 was linked to reduced expression of OAS3, an interferon-stimulated gene implicated in innate immune response and antiviral defense^49^. The decreased expression of OAS3 in T2D may impair proper immune signaling and contribute to sustained low-grade inflammation, a recognized contributor to insulin resistance and islet dysfunction^50,51^. Recent studies have also highlighted the role of other HLA class I polymorphisms in T2D susceptibility^52^. For example, Jan et al. (2021) identified a strong association between the HLA-B variant rs2308655 and T2D in the Pashtun population, with the minor C allele conferring over a sevenfold increased risk^53^. Although our results focused on HLA-A, these findings underscore the broader relevance of HLA class I gene variants in modulating immune responses and contributing to T2D pathophysiology. These results align with published evidence that links HLA polymorphisms and OAS3 dysregulation to immune system imbalances and chronic inflammation in T2D^54,55^. Importantly, these genes participate in an interlinked network through processes such as lipid-induced β-cell stress, metabolically driven inflammation, and impaired insulin signaling, suggesting that the associated SNPs may not act in isolation but as part of interconnected regulatory networks.

## Conclusions

Multi-omic integration enables the identification of functional variants, enhances mechanistic understanding, and informs future polygenic risk models and therapeutic strategies.

By extending our analysis to include GWAS variants with p-values ≤ 0.0001 (below the conventional genome-wide significance threshold), we revealed sub-threshold variants that are functionally active to exert measurable effects on gene and protein expression. This approach highlights the utility of a more inclusive threshold in uncovering biologically meaningful signals that may otherwise be overlooked in traditional GWAS pipelines.

These converging patterns highlight the value of multi-omic integration for identifying causal mechanisms and inform the development of functionally grounded biomarkers. Future research should explore the predictive value of these molecular changes for disease onset and progression and assess their utility in enhancing polygenic risk models for precision prevention and therapeutic targeting.

## ACKNOWLEDGEMENTS

This work was supported by the Chan Zuckerberg Initiative’s Foundation for Accelerate Precision Health Program to Advance Genomics Research at Meharry Medical College (CZIF2022-007043); by the National Institute of Health under award numbers EY033264, GM151274, UC2MD019626; and by the Veterans Affairs Office of Research and Development under award number CX002780. Dr Ayo Priscille Doumatey, National Institutes of Health – National Human Genome Research Institutes.

